# Transcriptome analysis of the mouse fetal and adult rete ovarii and surrounding tissues

**DOI:** 10.1101/2023.11.06.565717

**Authors:** Dilara N. Anbarci, Rebecca O’Rourke, Yu Xiang, Derek T. Peters, Blanche Capel, Jennifer McKey

## Abstract

The rete ovarii (RO) is an epithelial structure that arises during fetal development in close proximity to the ovary and persists throughout adulthood in mice. However, the functional significance of the RO remains elusive, and it has been absent from recent discussions of female reproductive anatomy. The RO comprises three distinct regions: the intraovarian rete (IOR) within the ovary, the extraovarian rete (EOR) in the periovarian tissue, and the connecting rete (CR) linking the EOR and IOR. We hypothesize that the RO plays a pivotal role in maintaining ovarian homeostasis and responding to physiological changes. To uncover the nature and function of RO cells, we conducted transcriptome analysis, encompassing bulk, single-cell, and nucleus-level sequencing of both fetal and adult RO tissues using the *Pax8-rtTA; Tre- H2B-GFP* mouse line, where all RO regions express nuclear GFP. This study presents three datasets, which highlight RO-specific gene expression signatures and reveal differences in gene expression across the three RO regions during development and in adulthood. The integration and rigorous validation of these datasets will advance our understanding of the RO’s roles in ovarian development, female maturation, and adult female fertility.

**Short narrative:** This study employs comprehensive bulk, single cell and single nucleus transcriptome analysis to uncover gene expression signatures of the fetal and adult rete ovarii (RO).

## Background and Summary

The rete ovarii (RO) is an epithelial structure that develops in close association with the ovary during fetal life and remains in adulthood in mammals (1,2). Despite the significant architecture of this ovarian appendage, and the fact that it is highly conserved in mammals (3–5), the function of the RO has not yet been determined, and it has disappeared from recent descriptions of female reproductive anatomy. The RO is the female homolog to the rete testis, thought to arise from the mesonephric tubules (1,6). The RO is divided into three regions; the intraovarian rete (IOR), which resides inside the ovary, the extraovarian rete (EOR) located in the periovarian tissue, and the connecting rete (CR), which links the EOR and IOR (6). Using the mouse as a model system we developed tissue-clearing and 3D-imaging methods using lightsheet microscopy (7) that allowed us to observe the RO in unprecedented detail (2). The bipotential rete structure first appears in both sexes as a PAX8+ population of cells at the interface between the dorsal aspect of the gonad and the mesonephros (8,9). This population of PAX8+ cells was recently shown to give rise to a subset of gonadal supporting cells in both sexes (9), and remains after gonadal sex determination as the IOR in females. The EOR begins to develop from the mesonephric tubules as a blind tubular epithelium that connects to the CR starting around E14.5 (2). In the adult, the IOR has regressed to a smaller population of cells within the ovary, while the EOR has significantly expanded into a single convoluted tubule ending in a blind distal dilated tip (2,10). The RO is in a unique location between the ovary and extraovarian milieu, where vascular and neuronal networks enter the ovary. We hypothesize that proximity to vascular and neuronal networks might allow the RO to sense homeostasis and convey information to the adult ovary. To better understand the nature and function of cells within the RO, we performed unbiased high-throughput transcriptome analysis of the RO and tissues that surround it. Here, we report three datasets that sequence the fetal and adult RO transcriptome in bulk, and at the single cell and single nucleus level. We use these datasets to identify RO-specific gene expression signatures, and to further characterize gene expression differences between the three regions of the RO during development and in the adult. To allow for accurate capture and enrichment of RO cells in our sequencing samples, we used the *Pax8-rtTA; Tre-H2B-GFP* mouse line, in which all regions of the RO express nuclear GFP at all stages. Figure 1 shows the expression domain of GFP in the fetal (Fig.1A, *E16.5*) and adult (Fig.1A, *2M*) mouse ovary, along with an illustration of the sample collection and analysis workflow for each dataset generated (Fig. 1B). These datasets complement recent publications that specifically focused on the intraovarian region of the RO in mouse and human (9,11,12), and fill a critical gap in knowledge by providing data for all three regions of the RO in both adult and fetal stages. Integration and careful validation of all these datasets will pave the way towards understanding the roles of the RO, and its function in ovary development, female maturation, and adult female fertility.

**Figure 1.**
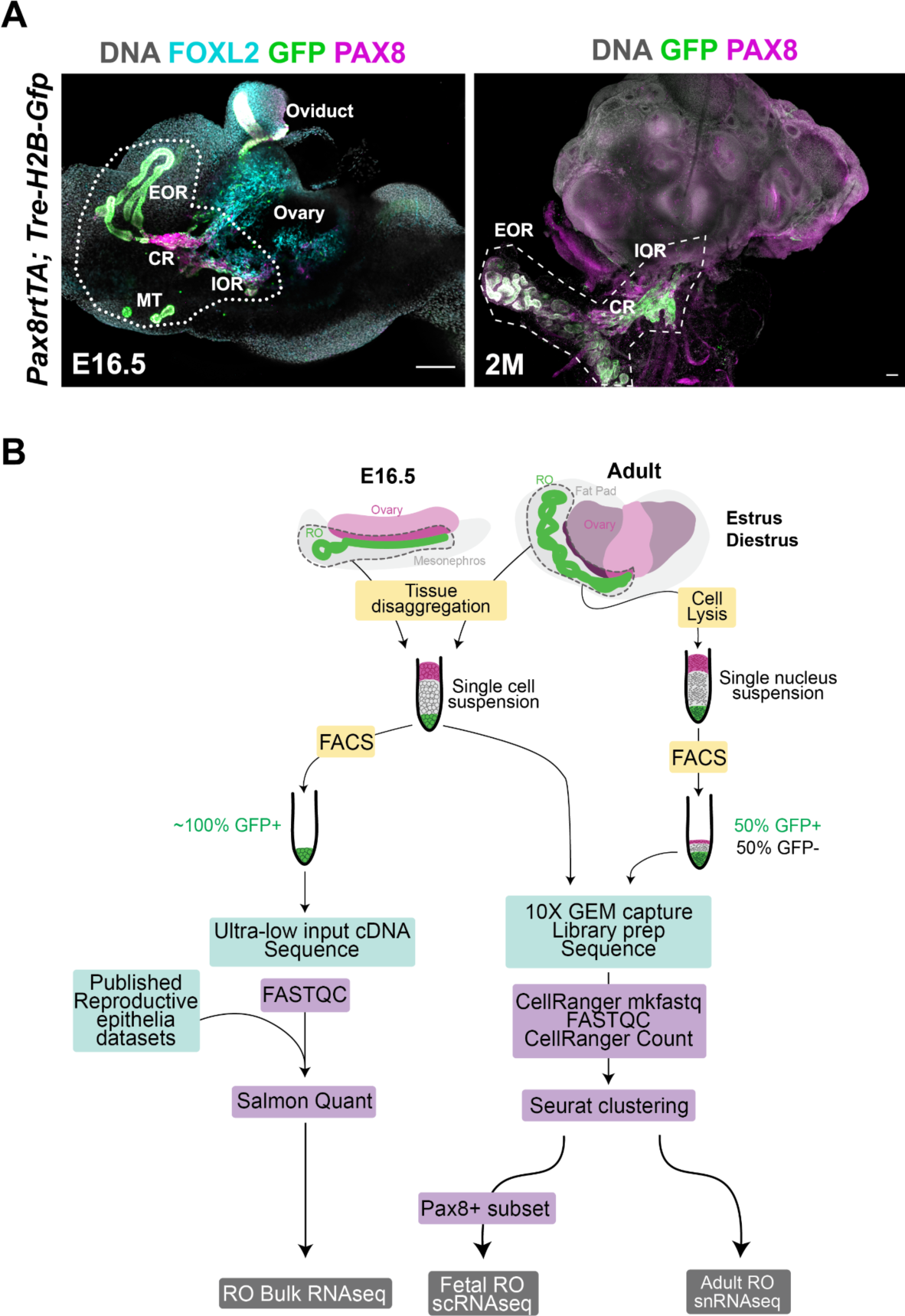
Overview of experimental design. (**A**) Maximum intensity projection of confocal Z-stack images of the ovary/mesonephros/oviduct complex of the E16.5 *Pax8rtTA; Tre-H2b-GFP* mouse embryo (*left panel*), and the ovary/fat pad complex of the 2-month-old *Pax8rtTA; Tre-H2b-GFP* mouse. The rete ovarii is labeled with PAX8 antibodies in *magenta* and GFP antibodies in *green* in both panels, and the ovary is labeled with FOXL2 antibodies in *cyan* in the E16.5 (left panel). Dotted lines around the rete ovarii estimate the amount of surrounding tissue collected for further analysis. *Scale bars, 100um*. EOR, extraovarian rete; CR, connecting rete; IOR, intraovarian rete; MT, mesonephric tubules. (**B**) Illustration of the workflow and analysis pipelines for the generation of each dataset.

## Methods

### Mice

All experiments were performed on *Pax8rtTA; Tre-H2b-Gfp* (PTG) female mice. The *Pax8-rtTA* (*B6.Cg-Tg(Pax8-rtTA2S*M2)1Koes/J;* RRID: IMSR_JAX:007176) and *Tre-H2BGFP (Tg(tetO-HIST1H2BJ/GFP)47Efu/J*; RRID:IMSR_JAX:005104) lines were previously described (13,14) and maintained on a mixed CD-1 / C57BL/6J background. To collect embryos at specific developmental stages, males were set up in timed matings with several females. Females were checked daily for the presence of a vaginal plug. Date of the plug was considered embryonic day 0.5. For adult timepoints, female carriers of the PTG alleles were weaned at 28 days and kept in single-sex housing until they reached 8-10 weeks old. Adult females for adult timepoints or pregnant dams for fetal timepoints were given a doxycycline diet of 625 mg/kg (Teklad Envigo TD.01306) 3 days prior to tissue collection to induce GFP expression in *Pax8*+ cells. For collection of adult tissue at specific stages of the estrous cycle, vaginal swabs were collected and vaginal cytology was examined to determine estrous stage based on previously described standard metrics (15). Female mice were estrous-tracked for a week prior to tissue collection to ensure they displayed typical cycling. Secondary validation of the estrous stage was performed by visual analysis of uterine swelling during tissue collection. Any mismatched observations resulted in tissue being omitted from further analysis. All mice were housed and handled in accordance with National Institutes of Health guidelines, in a barrier facility maintained at a temperature of 22 ± 0.1 °C, 30–70% humidity, within individually ventilated cages (Allentown; PNC75JU160SPCD3), and a controlled 12h on / 12h off light cycle. All experiments were conducted with the approval of the Duke University Medical Center Institutional Animal Care and Use Committee (IACUC protocol # A089- 20-04 9N).

### Whole-mount Immunostaining for confocal imaging

E16.5 ovary/mesonephros/Müllerian duct complexes and adult ovary/fat pad complexes were dissected in PBS-/- and fixed for 30 min (fetal) or 1 hour (adult) at room temperature in 4%PFA/PBS. Following two 15min PBS washes, samples were gradually dehydrated in MeOH dilutions (25% MeOH/PBS; 50% MeOH/PBS; 75% MeOH/PBS; 100% MeOH) for 15 min (fetal) or 30 min (adult) each at room temperature. Samples were stored at -20°C in 100% MeOH until required for staining. Before staining, samples were gradually rehydrated into PBS through 10min (fetal) or 20min (adult) washes in a reverse methanol gradient (75% MeOH/PBS; 50% MeOH/PBS; 25% MeOH/PBS), and transferred to PBS 0.1% Triton X-100 for 30min. Samples were then transferred to blocking solution (PBS; 1% Triton X-100; 10% horse serum) for 1h, and incubated at 4°C overnight (3 nights for adult) in primary antibodies diluted in blocking solution (chicken anti-GFP, 1:1000, RRID AB_300798; rabbit anti PAX8, 1:500, RRID AB_2236705; goat anti FOXL2, 1:250, RRID AB_2106188). The next day, samples were washed three times for 30min in PBS 0.1% Triton X-100 and incubated at 4°C overnight in secondary antibodies and Hoechst vital dye diluted 1:500 in blocking solution (AF647 Donkey anti rabbit, RRID AB_2492288; AF488 donkey anti chicken, RRID AB_2340375; Cy Donkey anti goat; RRID AB_2307351). On day 3, samples were washed twice for 15 min in PBS 0.1% Triton X-100 and transferred to PBS at 4°C until ready to mount for confocal imaging (1h - 48h).

Confocal images were captured in the longitudinal plane on a Zeiss LSM880 confocal microscope and the affiliated Zen software (Carl Zeiss, Inc., Germany) using a 10X objective.

### Fetal tissue collection

Embryonic day (E)16.5 fetuses were harvested from pregnant *Tre-H2BGFP^tg/tg^*dams crossed to a *Pax8rtTA^tg/+^* male. Embryos positive for both alleles were identified by the presence of green fluorescence in the urogenital epithelia. Male fetuses were discarded, and ovary/mesonephros/oviduct complexes from females were dissected in RNase-free PBS-/-, with as little removal of surrounding tissue as possible to ensure retention and survival of RO cells. The GFP signal was then used to perform close dissection of the RO and surrounding ovarian capsule, and the oviduct was discarded. To ensure capture of the IOR and CR, only the dorso-lateral-most portion of the ovary was conserved. The samples were kept on ice until all fetuses were dissected.

### Adult tissue collection

Ovaries and periovarian fat pads (in which the adult RO is embedded) were harvested from *Pax8rtTA*; *Tre-H2BGFP^tg/tg^* adult females. The RO was located within the fat pad by green fluorescence and closely dissected in RNase-free PBS-/-, with removal of as much surrounding tissue as possible to ensure enrichment of RO cells. To ensure capture of the IOR and CR, the dorso- lateral-most portion of the ovary was conserved, and the rest of the ovary was discarded. The samples were kept on ice until all samples were dissected. For single-nucleus RNA sequencing (snRNAseq) analysis of the adult RO, samples were collected from 5 mice in early diestrus and 5 mice in estrus (total of 10 ROs in each group), snap frozen in liquid nitrogen, and stored at -80°C until further processing.

### Bulk RNA sequencing

For each timepoint (E16.5 and 2month), three biological replicates were processed as follows. Pooled ROs (4-5 ROs for each replicate) were washed in RNase-free PBS-/- and incubated at 37°C in fresh 1X TrypLE (Thermo Fisher Scientific, catalog #12563029) for 20 minutes.

TrypLE was then aspirated off, and samples were resuspended in chilled RNase-free PBS with 3% bovine serum albumin (BSA) and pipetted gently up and down for about 2 min to disaggregate cells. Cells were pipetted through a 0.32μm cell strainer into FAC-sorting filter tubes (Corning Falcon cat # 352235). Single-cell suspensions were then FAC-sorted at the Duke Flow Cytometry Shared Resource core on a B-C Astrios Sorter, based on presence or absence of GFP. Cells were sorted into PBS, pelleted, and resuspended in 10.5ul Clontech Lysis buffer (Takara Bio Inc. Cat # 635013). Lysed samples were stored at -80°C until transferred to the Duke Center for Genomic and Computational Biology Sequencing and Genomic Technologies core facility for cDNA library preparation and next- generation sequencing. RNA quality control was measured on a TapeStation (Agilent Technologies) using High Sensitivity RNA ScreenTape (Agilent Technologies). All samples passed quality control.

Sample libraries were prepared with SMART-Seq v4 Ultra Low Input RNA Kit (Takara Clontech Kit Cat# 63488). Libraries were sequenced on a NovaSeq 6000 (Illumina) as 50bp paired-end reads (∼46 M reads/sample). Quality control was performed using FastQC (v 0.11.8). Reads were mapped to the GRCm39 mouse genome using Salmon with default settings (16). Mapped reads were annotated using the GRCm39 Ensembl Mus musculus gene annotation reference (release 110). Read abundance from Salmon (TPM) values were used for downstream quality control and gene expression analysis. For comparison to other reproductive tissues, FASTQs from published bulk RNAseq datasets were obtained from E16.5 ovary (17) (16.5dpc rep 1,2,3 from GSE117590), adult ovary (18) (control 1, 2, 3 from GSE101906), adult ovarian surface epithelium and adult oviduct (19) (Normal OSE and normal FTE reps 1,2,3 from GSE125016). Each FASTQ was reanalyzed using the same Salmon pipeline as for our RO datasets for accurate gene expression comparison.

### Sample preparation for single cell RNA sequencing of fetal RO

28 ROs at E16.5 and E17.5 were collected as described above, pooled and incubated for 12 min in 450ul Trypsin 0.05% with 50ul 2.5% Collagenase Type IV at 37°C. 500ul of chilled PBS 0.3% BSA were then added for mechanical disaggregation by pipetting gently up and down for about 2 min. Some tissue chunks were still present, thus the sample was incubated for another 6 min in 100ul Trypsin 0.05% with 50ul 2.5% Collagenase Type IV at 37°C. The sample was then pelleted by gentle centrifugation (5 min 500 x g 4°C) and resuspended in 500ul Red Blood Cell lysis buffer (eBioscience, cat #00-4333-57) and incubated for 3 min at room temperature. Cells were pelleted by gentle centrifugation and resuspended in 120ul PBS 0.3% BSA before passing through a 0.32um cell strainer into a clean 1.5mL tube. 10ul of the cell suspension was collected for viability assessment using Trypan blue and a hemocytometer. Manual counts determined that cell viability was 87.3%, with 49,500 live cells (∼400 cells / ul).

### Sample preparation for single nucleus RNA sequencing of adult RO

Pooled snap frozen ROs at diestrus (N=10) and estrus (N=10) were further separated into two subgroups for gentle (N=5) and harsh (N=5) dissociation to ensure sensitive and highly adherent cell types were present in the final single nucleus suspension. Samples were then processed independently as described in the demonstrated protocol for nuclei isolation provided by 10X Genomics (https://assets.ctfassets.net/an68im79xiti/6x4KMzpIgPgkje01sR1Xgr/9cfb7d859985e5c479aec4e0e501f903/CG000124_Demonstrated_Protocol_Nuclei_isolation_RevE.pdf). Briefly, frozen samples were placed in a dounce with 1mL of cell lysis buffer (10mM Tris-HCl pH 7.4; 10mM NaCl; 3mM MgCl2; 0.1%NP40) and left to incubate for 2 min on ice. Another 1.5mL of Cell Lysis Buffer was added, tissue was gently homogenized using the dounce, and left to incubate for another 3 min on ice. Finally, another 3mL lysis buffer was added and tissue was homogenized. For the light dissociation subgroups, tissue was gently homogenized, and tissue clumps remained in the solution, while for the harsh dissociation subgroups, the tissue was homogenized until no visible tissue clumps were left. After dissociation, nucleus suspensions were passed through a 70um cell strainer into a 50mL conical tube, and filtered again through 40um filter tips (Millipore Sigma Flowmi® Cell Strainers; BAH136800040) into a 15ml conical tube. Nucleus suspensions were then pelleted by gentle centrifugation (5 min, 500 x g, 4°C), supernatants were removed, and nuclei pellets were resuspended in 1mL chilled wash buffer (PBS 1%BSA, 0.2U/ul RNase out, Thermo Fisher cat #0777019). Samples were transferred to 1.5mL tubes and pelleted by gentle centrifugation, supernatants were removed, and pellets were resuspended in 1mL chilled wash buffer and transferred to FACS filter tubes (Corning Falcon cat # 352235). NucRed live 647 ready probe (Thermo Fisher Scientific, cat # R37106) was added to each tube 30 min prior to FAC- sorting to label live nuclei. Single nucleus suspensions were FAC-sorted at the Duke Flow Cytometry Shared Resource core on a B-C Astrios Sorter. NucRed-negative cells were discarded and NucRed- positive cells were sorted based on presence or absence of GFP. Our aim was to collect 50% GFP+ (putative RO) and 50% GFP- cells. Final nucleus counts were Estrus: 18054 GFP+ and 16619 GFP-; diestrus: 18337 GFP+; 18337 GFP-. GFP+ and GFP- nuclei for each stage were sorted into a single tube containing 700ul PBS, 1% BSA, pelleted by gentle centrifugation, and resuspended in 35ul PBS, 0.05% BSA.

### 10X Chromium Single Cell / Nucleus Capture, Library Preparation and Sequencing

To capture, label, and generate cDNA libraries of individual cells and nuclei, the 10X genomics Chromium Single Cell 3’ Library and Gel Bead Kit v3 following the 10X Genomics User Guide (https://assets.ctfassets.net/an68im79xiti/4tjk4KvXzTWgTs8f3tvUjq/2259891d68c53693e753e1b45e42de2d/CG000183_ChromiumSingleCell3v3_UG_Rev_C.pdf) was used. Briefly, the single cell / nucleus suspensions, RT-PCR master mix, gel beads and partitioning oil were loaded into a Single Cell A Chip 10X genomics chip, placed into the Chromium controller, and the Chromium single cell A program was run to generate GEMs (Gel Bead-In-EMulsion) that contain RT-PCR enzymes, cell lysates and primers for sequencing, barcoding, and poly-DT sequences. GEMs were then transferred to PCR tubes and the RT-PCR reaction was run to generate barcoded single-cell identified cDNA. Barcoded cDNA was used to make sequencing libraries. Sequencing was performed on an Illumina NovaSeq 6000 S-Prim using paired end 150 cycles 2x150 reads by the Duke Center for Genomic and Computational Biology Sequencing and Genomic Technologies core facility.

### 10X single cell and single nucleus RNAseq data analysis

The 10X Genomics Cellranger (v3.1.0) mkfastq software was used for FASTQ generation, quality control was performed using FastQC (v 0.11.8), and CellRanger Count was used for alignment, filtering, barcode counting, and UMI counting of the single cell/nuclei FASTQs. For the snRNAseq data, the argument “include-introns true” was added to account for nuclear RNA. Seurat (v4.3.0.1) (20,21) was used for cluster analysis using R (v4.1.2) in RStudio software (2023.06.1+524). The E16.5 scRNAseq dataset was filtered to remove cells with <200 genes, >7500 genes or percent mitochondrial genes >10% and clusters were found using FindNeighbors(reduction=’pca’, dims=1:42) and FindClusters(resolution=0.8). Sub-clusters for *Pax8*+ (PxPos object) cells were found by subsetting the data based on c(‘*Pax8’*)>1 and standard seurat analysis of the PxPos dataset with FindNeighbors(reduction=’pca’, dims=1:9) and FindClusters(resolution=0.6) The snRNAseq data from adult estrus and diestrus samples were merged and integrated using the FindIntegrationAnchors and IntegrateData functions of Seurat, and the integrated dataset was filtered to remove cells with <200 genes, >7500 genes or percent mitochondrial genes >10% and clusters were found using FindNeighbors(reduction=’pca’, dims=1:30) and FindClusters(resolution=0.5).

### Data Records

The sequencing data from this study have been uploaded to the National Center for Biotechnology Information (NCBI) Gene Expression Omnibus with accession ID GSE244849. This includes 6 raw.fastq.gz files for E16.5 and adult bulk RNAseq data; 3 raw.fastq.gz files for the E16.5 scRNAseq sample, and 3 raw.fastq.gz files for the adult snRNAseq data. We also provide the output from Salmon transcript quantification as *quant.sf* files with transcript abundance values for each replicate of our fetal and adult RO bulk RNAseq, as well as the reanalyzed E16.5 ovary, adult ovary, adult OSE, and adult oviduct datasets. In addition, we provide the *filtered_feature_bc_matrix* output folders from CellRanger Count for the E16.5 scRNAseq and adult snRNAseq datasets. All code generated for the analyses presented here is available on the McKey Lab GitHub at https://github.com/McKeyLab/RODatasets. In our GitHub repository, we also provide a .rmd (ROGeneTest.rmd) file to query all the datasets for expression of a specific gene. The output from this file includes a bar plot of TPMs from the bulk RO and reproductive epithelia analyses, and feature plots and violin plots for the fetal scRNAseq and the adult snRNAseq, for the full datasets and the *Pax8*+ subsets. All datasets can also be queried on our shiny app at https://cuanschutz-devbio.shinyapps.io/McKey_rete_ovarii_shiny/.

## Technical Validation

### Experimental overview

We set out to sequence the transcriptome of the RO at E16.5 and in the 2-3 month-old adult. To facilitate the capture of RO cells, we took advantage of the *Pax8rtTA; Tre- H2b-GFP* transgenic line, in which all regions of the RO express GFP during fetal development and in the adult (Fig. 1A). We collected samples of the RO and surrounding tissue from these mice and processed them for either Bulk RNAseq (E16.5 and adult), single-cell RNAseq (E16.5), and single-nucleus RNAseq (adult) (Fig. 1B). Standard pipelines were used in each case. For BulkRNAseq, we collected GFP+ cells using FACS analysis. Since the RO represents so few cells, we used the Ultra-Low input SMARTseq kit provided by Takara Clontech, to prepare cDNA libraries (Fig. 2B, *left*). For the E16.5 single-cell sample, we did not FAC-sort the cells, but rather collected the entire ovarian capsule for single-cell capture using 10X Genomics Chromium (Fig. 2B, *middle*). For the adult samples at Estrus and Diestrus, we learned from our E16.5 experience that it would be better to enrich the sample for RO cells and chose to FAC-sort approximately 50%GFP+ and 50%GFP- nuclei for snRNA sequencing. All cDNA libraries were sequenced using a NovaSeq 6000, resulting FASTQ read files were quality checked using FASTQC software and standard downstream analysis was performed. For Bulk RNAseq, transcript abundance and differential gene expression were obtained using Salmon and DESeq2, while 10X CellRanger and Seurat software were used to analyze single cell and single nucleus datasets (Fig. 2B).

**Figure 2.**
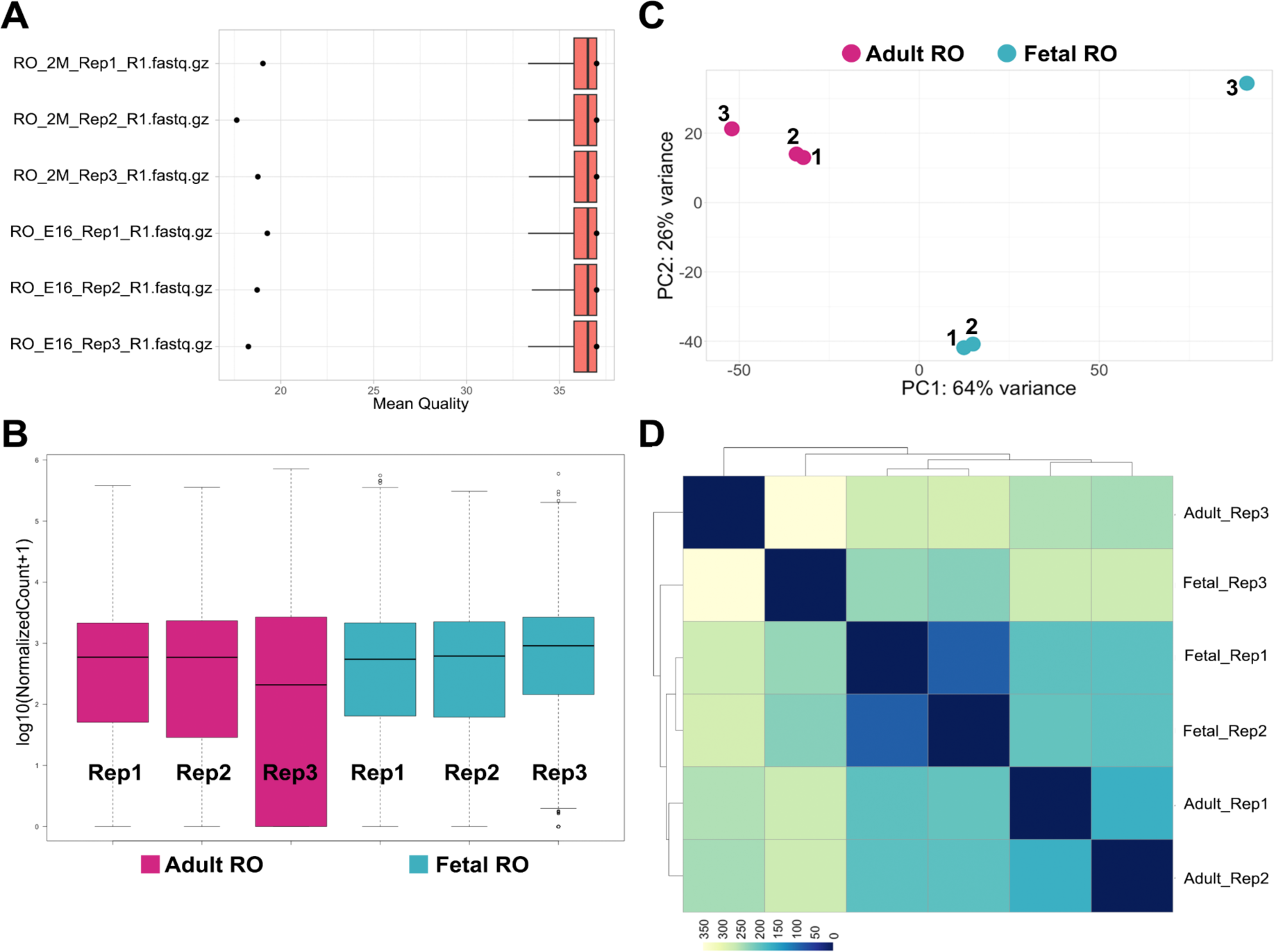
Technical validation of RO Bulk RNAseq datasets. (**A**) Boxplot illustrating normalized counts for each sample on the logarithmic scale. Adult RO replicates are depicted in *cherry,* fetal RO replicates in *teal*, feta; ovary. Note all samples follow a similar distribution except for Adult_RO rep3, which appears to be an outlier. (**C**) Scatter plot illustrating Principal Component Analysis (PCA) of Adult and fetal RO datasets. Each dot represents a single replicate for each timepoint (*cherry*, adult RO; *teal*, fetal RO). In each case, replicates 1 and 2 cluster more closely than replicate 3. In particular, fetal RO rep3 appears to be a true outlier. (**D**) Heatmap showing sample distance analysis for all adult RO and fetal RO replicates. Replicates 1 and 2 for each stage share more similarities than replicates 3, as illustrated by the deeper blue hues in the bottom right third of the heatmap. Rep1, replicate 1; Rep2, replicate 2; Rep3, replicate 3.

### Bulk RNAseq of the fetal and adult rete ovarii

We first performed bulk RNA sequencing of FAC- sorted GFP+ cells for three biological replicate samples of pooled ROs (4-5 per pool) from *Pax8-rtTA; TreH2bGFP* females at E16.5 and 2 months (Table 1). Paired-end 50bp sequencing runs performed within the standard quality range, and we achieved 53-84M read pairs for each sample (Table 1). We used FASTQC (22) and Rqc (23) for FASTQ quality control, and found that the Mean Read quality passed the standard quality threshold of 35 for all samples (Fig. 2A). We then used Salmon software (16) to map reads to the mouse genome (build GRCm39) and quantify transcript expression. Salmon mapping rate ranged from 68% to 87% across all samples (Table 1). The output from Salmon was brought into RStudio software for count normalization (Fig. 2B). These results showed that the normalized counts for all samples fell in the same range, except for the adult replicate 3, which had on average lower counts. Principal component analysis (PCA, Fig. 2C), and sample distance analysis (Fig. 2D) showed that Fetal RO replicate 3 was a major outlier in the experiment, and that Adult RO replicate 3 was somewhat of an outlier.

**Table 1.**
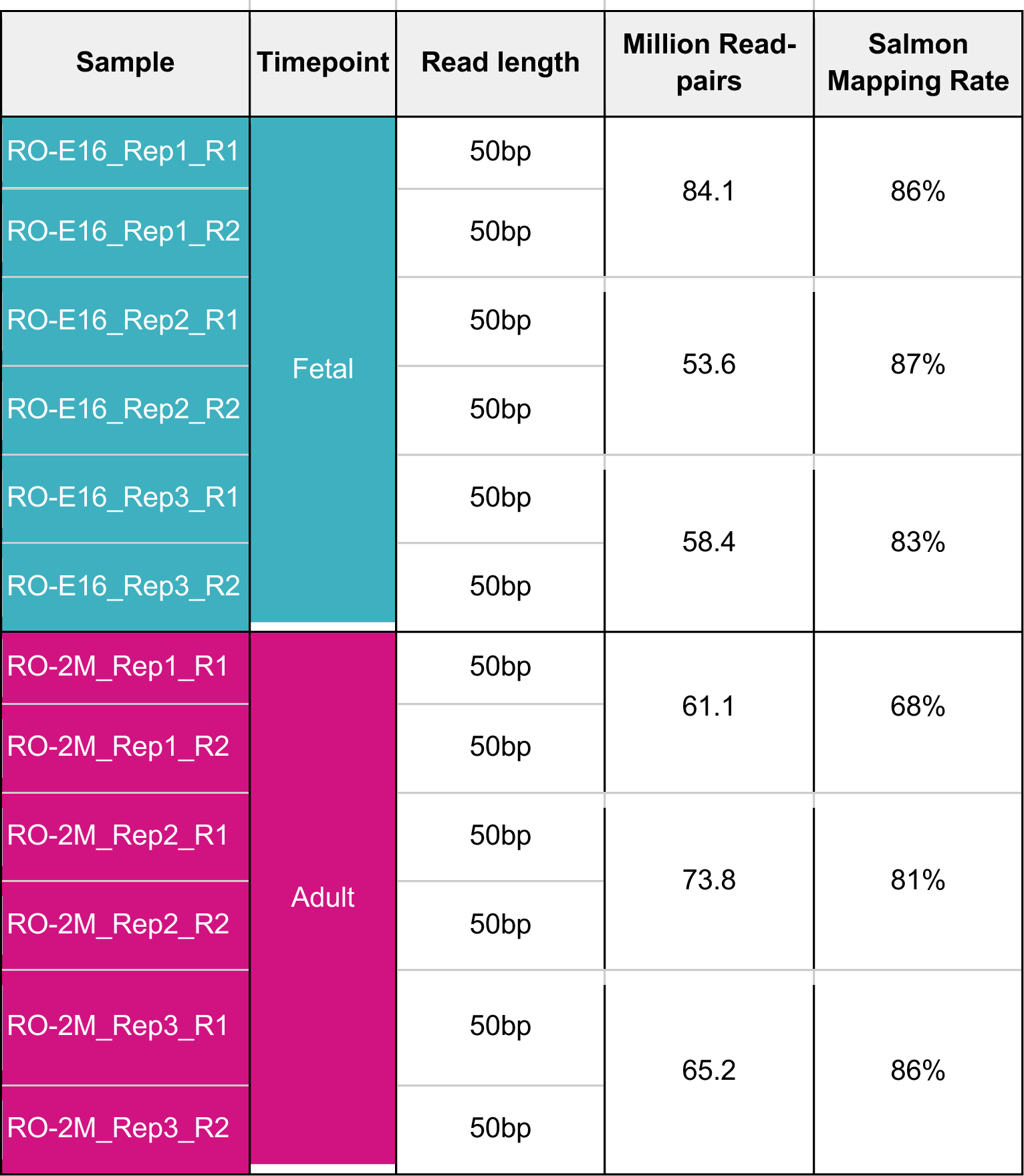
Rete ovarii Bulk RNA sequencing run Information.

| Sample | Timepoint | Read length | Million Read-pairs | Salmon Mapping Rate |
| --- | --- | --- | --- | --- |
| RO-E16_Rep1_R1 | Fetal | 50bp | 84.1 | 86% |
| RO-E16_Rep1_R2 |  | 50bp |  |  |
| RO-E16_Rep2_R1 |  | 50bp | 53.6 | 87% |
| RO-E16_Rep2_R2 |  | 50bp |  |  |
| RO-E16_Rep3_R1 |  | 50bp | 58.4 | 83% |
| RO-E16_Rep3_R2 |  | 50bp |  |  |
| RO-2M_Rep1_R1 | Adult | 50bp | 61.1 | 68% |
| RO-2M_Rep1_R2 |  | 50bp |  |  |
| RO-2M_Rep2_R1 |  | 50bp | 73.8 | 81% |
| RO-2M_Rep2_R2 |  | 50bp |  |  |
| RO-2M_Rep3_R1 |  | 50bp | 65.2 | 86% |
| RO-2M_Rep3_R2 |  | 50bp |  |  |

### Comparison between the bulk transcriptome of the RO and other reproductive epithelia

To further analyze the specific gene expression signatures of the RO, we accessed and re-analyzed published bulk RNAseq datasets of other reproductive epithelium and the ovary. For this, we chose datasets from E16.5 ovary (17), adult ovary (18), adult ovarian surface epithelium and adult oviduct (19), as these were all expected to be the most closely related to the RO, and differential gene expression was likely to point us towards RO-specific genes. Each dataset had three replicates, which were analyzed alongside our RO datasets using Salmon quantification (Fig. 3). Log10-normalized counts were in the same range for all samples (except, as noted before, Adult RO replicate 3) (Fig. 3A). PCA and sample distance analysis showed that all replicates from each cell type clustered together (Fig. 3B, C). Interestingly, the fetal and adult RO (Fig. 3B, *teal* and *cherry*, respectively) clustered close to each other, but further away from the other reproductive epithelia (Fig. 3B). This could be due to a batch bias in the data generation, since they were sequenced separately from the other samples. On the other hand, the adult ovary and adult ovarian surface epithelium (OSE) clustered closely despite being generated in two different studies (Fig. 3B, *yellow* and *orange*, respectively). This was to be expected, since the ovary dataset likely included OSE. In addition, these tissues are very closely related, and the OSE is a source of ovarian cells (24,25). The fetal ovary and adult oviduct (Fig. 3B, *green* and *purple*, respectively) were the furthest from the other tissues (Fig. 3B, C), with the fetal ovary clustering almost equidistant to the fetal RO and to the adult ovary. Intriguingly, the adult RO samples appeared closer to the adult OSE than to the adult ovary.

**Figure 3.**
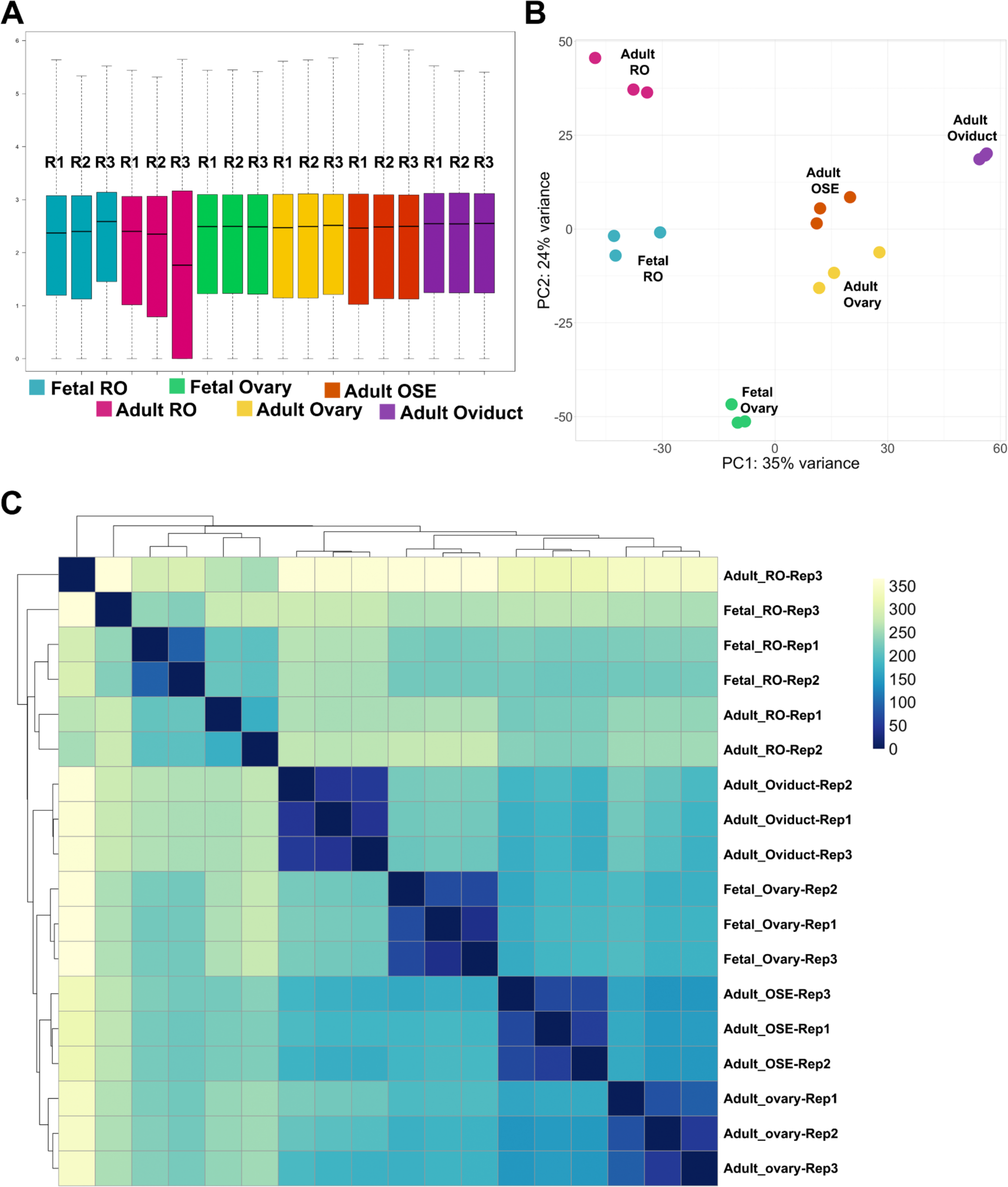
Technical validation of reproductive epithelia Bulk RNAseq datasets. (**A**) Boxplot illustrating normalized counts for each sample on the logarithmic scale. Adult RO replicates are depicted in *cherry,* fetal RO replicates in *teal*, fetal ovary replicates in *green*, adult ovary replicates in *yellow*, adult ovarian surface epithelium (OSE) replicates in *orange*, and adult oviduct (ovi) samples in *purple*. Note all samples follow a similar distribution except for Adult_RO rep3, which appears to be an outlier. (**B**) Scatter plot illustrating Principal Component Analysis (PCA) of all reproductive epithelia datasets. Each dot represents a single replicate for each timepoint (*cherry*, adult RO; *teal*, fetal RO; *green*, fetal ovary; *yellow*, adult ovary; *orange*, adult OSE; *purple*, adult oviduct). In each case, all 3 replicates for a tissue cluster together more closely than with any other tissue. (**C**) Heatmap showing sample distance analysis for all adult RO and fetal RO replicates. Rep1, replicate 1; Rep2, replicate 2; Rep3, replicate 3.

We next performed differential gene expression to begin highlighting the differences between the RO and other reproductive epithelia. For this, we used the DESeq2 package in R (26). We performed pairwise differential gene expression (DGE) analysis between the Adult and fetal RO (Fig. 4A), and between the fetal or adult RO and the fetal ovary, adult ovary, adult OSE and adult oviduct (Fig. 4B). Top enriched genes in the adult RO vs. the fetal RO included *Fam180a, Efemp1, Gbp2, Igfbp6 and Lgr5*, while genes enriched in the fetal RO (negative L2FC) included *Ifitm1, Alpk1, Nell1 and Tex12* (Fig. 4A). From the comparison with other epithelia, genes that appear specific to the fetal RO compared to the fetal ovary include *Comt, Klkb1, Sass6, Zcchc17* (Fig. 4B, *top left*), and genes enriched in the adult RO in all comparisons include *Brms1l*, *Sass6*, *Comt* and *Bmt2* (Fig. 4B, *top right and bottom*). *Sass6* and *Comt* thus appear as potential markers of the RO at both stages. Further spatial validation using RNA visualization methods are required to test whether these are true markers of the RO.

**Figure 4.**
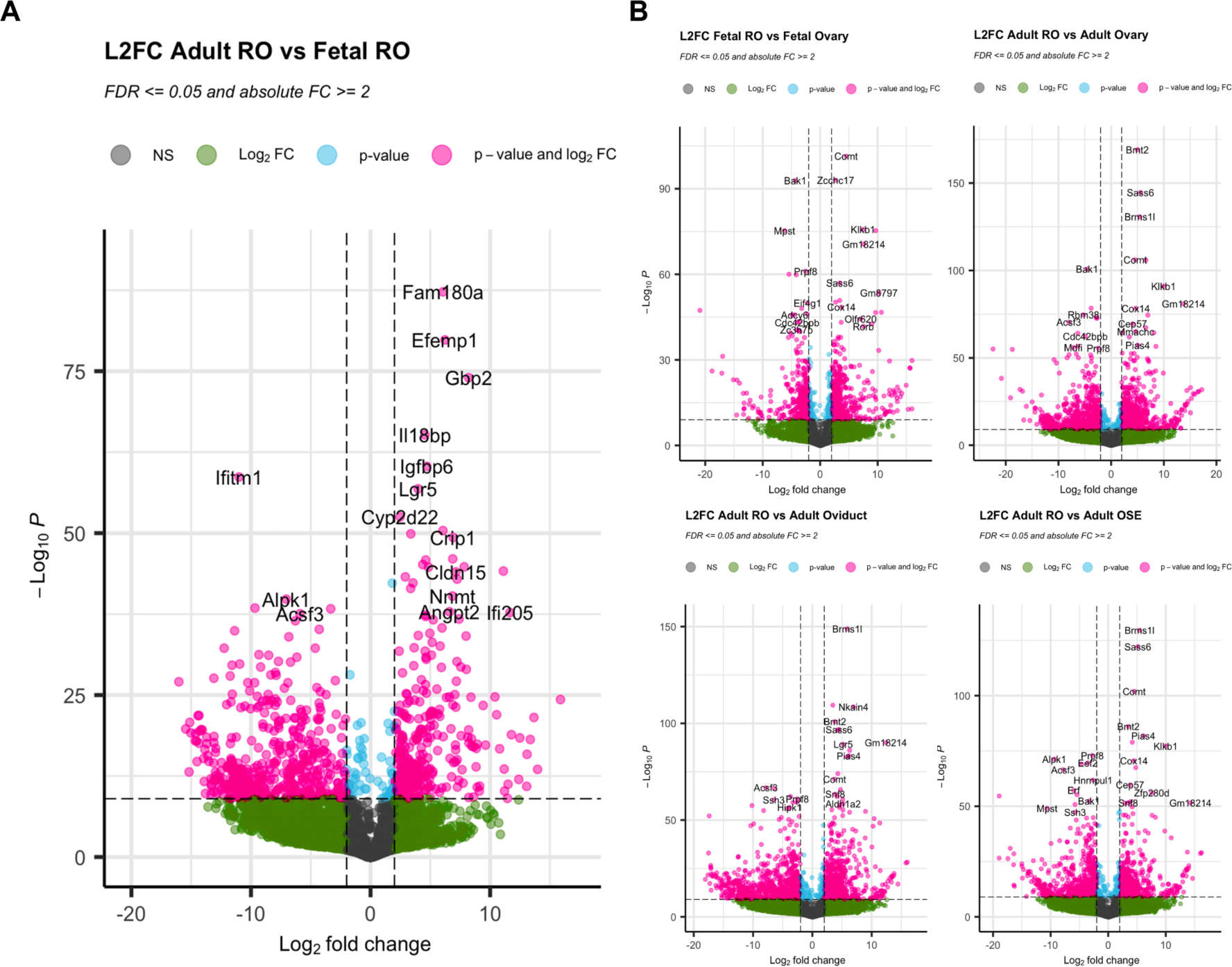
Differential gene expression analysis. (**A, B**) Volcano plots illustrating differential gene expression analysis comparing the adult RO to the fetal RO (**A**), and the RO vs other reproductive epithelia (fetal RO vs. fetal ovary, *top left*; adult RO vs. adult ovary, *top right*; adult RO vs adult oviduct, *bottom left*; adult RO vs. adult OSE, *bottom right*). Plots represent Log2 Fold change (L2FC) on the x- axis and -Log10(P Value) on the y axis. Gray dots are genes that are not significantly differentially expressed (NS); green dots are genes that display at least 2 or -2 L2FC (vertical dashed lines); blue dots represent genes with a significant L2FC (-Log10(P)>10, dashed horizontal line); pink dots with gene symbols are genes that are are significantly enriched (>2 L2FC) or downregulated (<-2 L2FC) in the first condition (**A**, adult RO; **B**, adult or fetal RO) compared to the second condition (**A**, fetal RO; **B**, other reproductive epithelium).

### Single-cell RNAseq of the fetal RO

To determine how the different regions of the RO are defined during development and begin to identify region-specific cellular functions, we next performed single-cell RNA sequencing of the RO. We collected ROs from E16.5 *Pax8rtTA; TreH2BGFP* females, dissecting as close to the RO as possible, but intentionally leaving some surrounding tissue from the ovarian capsule and mesonephric tubules. We hypothesized that this would allow us to identify markers of other poorly described components of the ovarian capsule, such as the mesovarium and metanephric mesenchyme. In addition, including this tissue meant we could interrogate the data for potential interactions between cells of the RO and surrounding tissue. We produced single cell suspensions and performed cell capture and barcoding using the 10X Genomics Chromium. After library preparation and sequencing, we performed QC on the FASTQ read files, studied the CellRanger QC output, and found that the reads passed QC thresholds (Table 2). We imported the outputs from CellRanger counts into R as a Seurat object, and began the standard Seurat single-cell analysis pipeline with the whole sample. Standard QC metrics for single cell data include the number of genes per cell (nFeature_RNA), which allows the distinction between lysed, empty cells (nFeature_RNA < 200) and cell doublets (nFeature_RNA > 7500), and the percentage of mitochondrial genes, which acts as a proxy for quantifying cell death (optimal percent.mt < 20) (Fig. 5A, B). Figure 5A shows that a majority of cells in our sample fall within the optimal quality range (Fig. 5A, *dotted rectangle*). Figure 5B shows violin plots for each QC metric and illustrates that most of our cells are within the optimal range (Fig. 5B). Nonetheless, we opted to filter out low quality cells (cells with <200 genes, >7500 genes or percent mitochondrial genes >10%) for further analysis, as recommended in the Seurat user guide. We then performed Seurat CLuster analysis and identified 22 clusters (Fig. 5C). Analysis of Pax8 expression in this dataset revealed that we had captured very few Pax8+ cells(Fig. 5D). Nonetheless, these cells mapped to distinct clusters. To analyze these cells further, we created a new dataset based on Pax8 expression (c(Pax8)>1), which we called PxPos. This subset was expected to contain only cells of the RO and the mesonephric tubules. Standard Seurat Analysis on this subset revealed 6 independent Pax8+ clusters (Fig. 5E). We used prior knowledge and the EnrichR gene ontology database to identify putative cell types for each cluster. *Foxl2* and *Nr5a1* are granulosa markers that are expressed in cells of the IOR (*Foxl2+* and *Nr5a1+*). (2,9).

**Figure 5.**
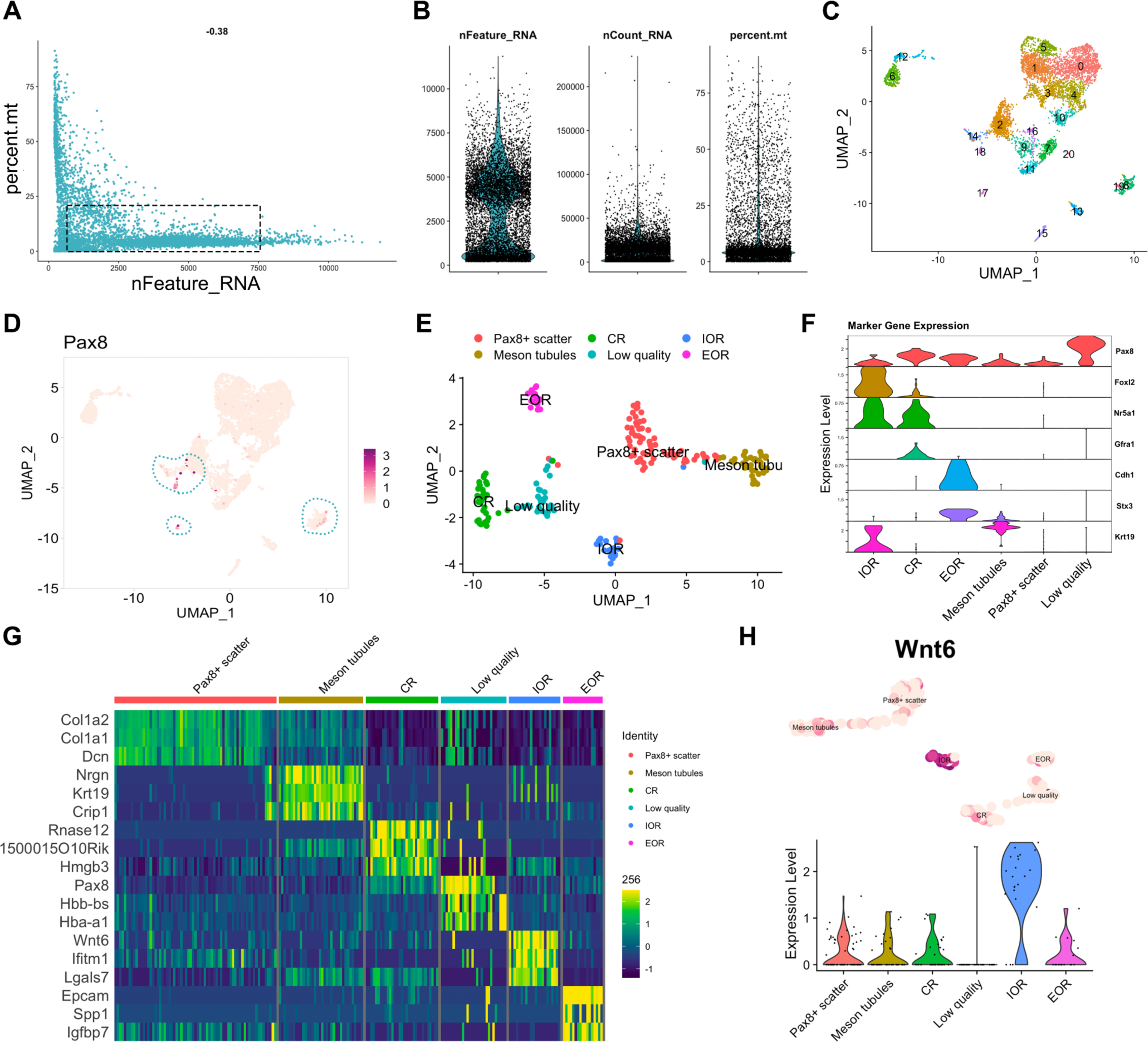
Technical validation of the E16.5 scRNAseq dataset. (**A, B**) Quality control of E16.5 single- cell data. (**A**) Scatter plot illustrating the distribution of the number of genes (nfeature_RNA) relative to the percentage of mitochondrial genes (percent. mito). Live cells of optimal quality have nFeature_RNA between 200 (empty cells) and 7500 (dublets), and have low expression of mitochondrial genes (used as a proxy for dying cells). The dotted rectangle represents the optimal quality threshold region. (**B**) Violin plots illustrate the number of genes (nfeature_RNA), unique molecular identifier (UMI) (nCount_RNA), and the percentage of mitochondrial genes (percent. mito). (**C**) Seurat UMAP plot showing cluster analysis of the whole single-cell sample at E16.5, using nDims = 42 and resolution = 0.8. 21 clusters were identified, numbered 0 to 21. (**D**) Seurat FeaturePlot showing expression of *Pax8* across the whole single-cell dataset. Darker dots represent higher *Pax8* expression levels. Dotted outlines highlight the clusters that express Pax8 (clusters # 8, 14, 17, 19). (**E**) Seurat UMAP plot showing cluster analysis of the subset of *Pax8*+ cells (c(*Pax8*)>1, object = *PxPos*), using nDims = 9 and resolution = 0.6. 6 clusters were found, and putative cell types for each cluster were identified using known gene markers and Gene Ontology analysis. (**F**) Seurat Stacked Violin plot showing expression of marker genes in each cluster of the *PxPos* subset dataset. (**G**) Heatmap illustrating expression differences for top 3 enriched genes from each cluster in the PxPos dataset. Dark blue hues depict low expression while green-yellow hues depict high expression. (**H**) Illustration of the “gene expression layout” code chunk in our R markdown file for E16.5 scRNAseq expression analysis. This code chunk produces Seurat violin and Feature plot for a given gene. The example chosen here is *Wnt6*, which appears enriched in the IOR cluster of the E16.5 PxPos dataset.

**Table 2.** Cell Ranger metrics on fetal RO scRNAseq and adult RO snRNAseq runs.

|  | E16.5 scRNA | 2M snRNA<br>Diestrus | 2M snRNA<br>Estrus |
| --- | --- | --- | --- |
| **CellRanger count** | 7,952 | 2,964 | 2,792 |
| **Mean Reads per Cell** | 125,706 | 164,427 | 172,023 |
| **Valid Barcodes** | 96.90% | 95.20% | 94.80% |
| **Valid UMIs** | 99.90% | 99.90% | 99.90% |
| **Reads Mapped to Genome** | 92.80% | 91.20% | 92.10% |
| **Reads Mapped Confidently to Genome** | 88.50% | 88.30% | 89.20% |
| **Reads Mapped Confidently to Intergenic Regions** | 4.50% | 4.50% | 4.40% |
| **Reads Mapped Confidently to Intronic Regions** | 23.60% | 60.70% | 59.20% |
| **Reads Mapped Confidently to Exonic Regions** | 60.40% | 23.10% | 25.70% |
| **Reads Mapped Confidently to Transcriptome** | 56.20% | 39.50% | 41.20% |
| **Reads Mapped Antisense to Gene** | 2.50% | 44.10% | 43.40% |

Cells of the CR express *Nr5a1+* and GFRa1, which is specific to the CR (Anbarci et al, 2023 -*preprint*). Thus we labeled the *Foxl2+/Nr5a1+/Gfra1-* cluster IOR, and the *Foxl2-/Nr5a1+/Gfra1+* cluster CR. We had previously found that at E16.5, E-Cadherin and STX3 are specifically expressed in the EOR (2)(Anbarci et al, 2023 -*preprint*), thus we called the *Cdh1*+/*Stx3*+ cluster EOR. Finally the remaining cluster was enriched in *Krt19*, which is not found in the EOR at this stage, but is present in the mesonephric tubules. Stx3 is generally a marker of tubular epithelia, thus we determined that the *Stx3*+/*Krt19*+ cluster likely represents mesonephric tubules remaining in the ovarian capsule (Fig. 5F). Using the *FindAllMarkers* function of Seurat, we identified the top 3 enriched genes for each cluster in the PxPos dataset. These included *Nrgn*, *Krt19* and *Crip1* as markers of the mesonephric tubule cluster; *RNase12*, *1500015O10Rik* and *Hmgb3* for the CR cluster; *Wnt6*, *Ifitm1* and *Lgals7* for the IOR cluster; and *Epcam*, *Spp1* and *Igfbp7* for the EOR cluster (Fig. 5G). Encouragingly, these genes were also found to be enriched in the bulk RNAseq dataset of the fetal RO. Spatial analysis of RNA expression using *in situ* hybridization will be required to validate the mapping of these clusters and these potential regional markers. The code used to perform these analyses and generate the panels in figure 5 is available on our GitHub page (https://github.com/McKeyLab/RODatasets). In the last chunk of these R markdown files, we provide the code to generate a FeaturePlot + Violin plot layout for any given gene. An example of one such layout is provided in figure 5H with *Wnt6*, a newly identified marker of the IOR.

### Single-nucleus RNAseq of the adult RO

We next sought to produce the single cell transcriptome of the adult RO. Having learned from the fetal time point that the number of RO cells is too low for optimal analysis, we chose to FAC-sort GFP+ cells from adult *Pax8rtTA; TreH2bGFP* females and mix GFP+ cells with GFP- cells in the final sample, to reach an optimal number of cells for 10X capture, and to allow for analysis of the interaction between cells of the adult RO and surrounding tissue. Our first disaggregation trials to generate single cell suspensions of the adult RO revealed that the tubular EOR is strongly adherent and very difficult to dissociate without lysing the cells. We thus turned to single-nucleus RNA sequencing, which is recommended for difficult tissues. Finally, we chose to sequence the RO from mice at estrus and diestrus, so that we can begin to interrogate whether gene expression in the RO changes during the estrous cycle. Similar to the fetal scRNAseq, single nuclei were captured using the 10X chromium, and cDNA libraries were prepared and sequenced. We began the analysis by performing QC on the FASTQ files and CellRanger outputs (Table 2), and both samples passed the QC thresholds. We then brought both datasets into Rstudio and merged them for analysis with Seurat. The QC metrics shown in the nFeature_RNA relative to percent.mt scatter plot (Fig. 6A) and violin plots (Fig. 6B) revealed that the majority of the cells in both samples were found within the optimal thresholds.

**Figure 6.**
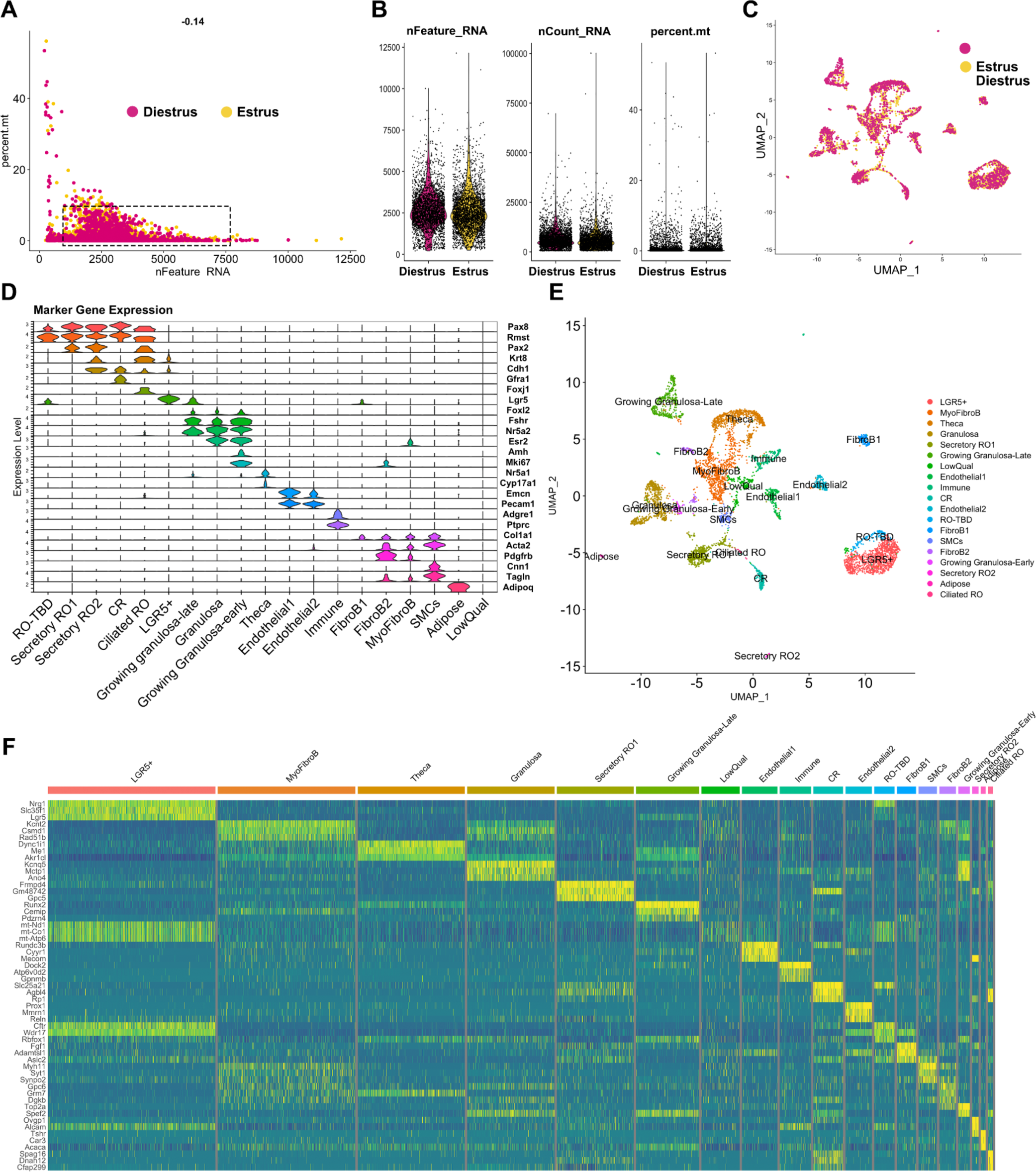
Technical validation of the adult snRNAseq dataset. (**A, B**) Quality control of Adult single-nucleus data at estrus (*yellow*) and diestrus (*cherry*). (**A**) Scatter plot illustrating the distribution of the number of genes (nfeature_RNA) relative to the percentage of mitochondrial genes (percent. mito). Live cells of optimal quality have nFeature_RNA between 200 (empty cells) and 7500 (dublets), and have low expression of mitochondrial genes (used as a proxy for dying cells). nFeature_RNA and percent.mt are lower here due to the sample being single nuclei. The dotted rectangle represents the optimal quality threshold region. (**B**) Violin plots illustrate the number of genes (nfeature_RNA), unique molecular identifier (UMI) (nCount_RNA), and the percentage of mitochondrial genes (percent. mito). (**C**) Seurat UMAP plot showing analysis of the integrated single-nucleus dataset from Estrus (*yellow*) and Diestrus (*cherry*) adult samples, illustrating high overlap between Estrus and Diestrus samples. (**D**) Seurat Stacked Violin plot showing expression of marker genes in each cluster of the *integrated* dataset. Putative cell types for each cluster were identified using known gene markers and Gene Ontology analysis. (**E**) Seurat UMAP plot showing cluster analysis of the single-nucleus integrated dataset, using nDims = 30 and resolution = 0.5. 19 clusters were identified, numbered 0 to 19. (**F**) Heatmap illustrating expression differences for top 3 enriched genes from each cluster in the Integrated snRNAseq dataset. Dark blue hues depict low expression while green-yellow hues depict high expression.

In this single-nucleus dataset, only RNAs found in the nucleus are represented. This explains the higher rate of intronic sequences and the higher rate of antisense reads identified in the single-nucleus dataset compared to the single-cell dataset of (Table 2). RNAs of mitochondrial genes are typically not present in the nucleus, thus the percent.mt was expected to be lower. In addition, because single-nucleus only captures nascent RNA in the nucleus, the number of RNAs (nCount_RNA) was also expected to be lower, as seen in the QC plots (Fig. 6A, B) (27). The number of genes and reads sequenced per cell was lower in the snRNA-seq compared to the scRNA-seq approach (Fig. 1C). After QC, the estrus and diestrus datasets were integrated in Seurat, and the UMAP of the integrated dataset showed that there was significant overlap between the two stages, with only a few clustered cells found in one stage but not the other (Fig. 6C).

To confirm that we had indeed captured the RO at estrus and diestrus, we investigated the expression of genes previously reported to change with the estrous cycle (28). *Star* and *Rgcc* were shown to be upregulated during estrus in granulosa and theca cells, and this is indeed what we found in our dataset (Fig. 7A). During diestrus, *Lhcgr* and *Inhba* are upregulated (28), and this was also true in our dataset (Fig. 7B). Thus, we conclude that our dataset does capture expression in the RO and surrounding cells at estrus and diestrus. Seurat clustering yielded 19 clusters.We attempted to identify cell types for each cluster using our prior knowledge and the EnrichR gene ontology database (29–31). The following markers were used to annotate the clusters: Epithelial markers (RO; OSE): *Krt8*; *Cdh1*, Epithelial progenitor marker (OSE, RO?): *Lgr5*; Ciliated epithelial marker: *Foxj1;* RO markers: *Pax8, Pax2, Gfra1, Rmst*; Granulosa cell markers: *Foxl2, Esr2;* Growing granulosa markers: *Amh, Nr5a2, Mki67;* Theca markers: *Nr5a1, Cyp17a1;* Immune cell markers: *Adgre; Ptprc;* Endothelial cell markers: *Pecam1, Emcn;* Adipose marker: *Adipoq;* Fibroblast / Myofibroblast markers: *Pdgfrb, Acta2, Col1a1;* Smooth muscle cell (SMC) markers: *Tagln, Cnn1* (Fig. 6D, E). Using the Seurat FindMarkers function, we identified the top 3 genes for each putative RO cluster as follows: Secretory RO1, *Frmpd4, Gm48742, Gpc5;* CR, *Slc25a21, Agbl4, Rp1*; RO-TBD, *Cftr, Wdr17, Rbfox1*; Secretory RO2, *Ovgp1, Alcam,Tshr*; Ciliated RO, *Spag16, Dnah12, Cfap299* (Fig. 6F). We are currently performing spatial validation for all the top genes, and hope this data will lead to exciting discoveries about the role and regulation of the RO.

**Figure 7.**
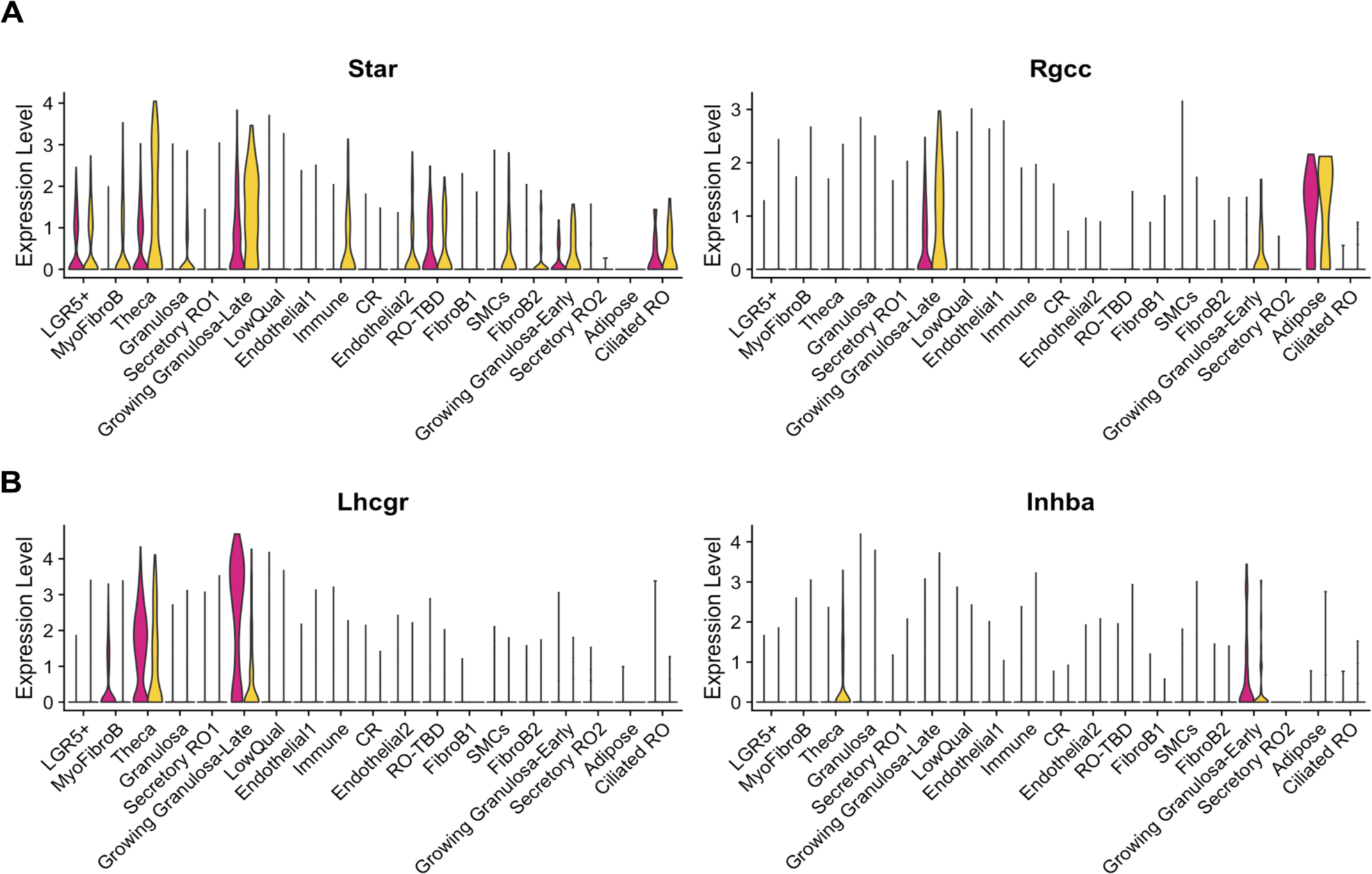
Validation of estrous staging for snRNAseq dataset. (**A, B**) Seurat violin plots illustrating expression of known cycling genes in the integrated snRNA dataset at estrus (*yellow*) and diestrus (*cherry*). Star and Rgcc are enriched in granulosa and theca cells at estrus (**A**), while Lhcgr and Inhba are enriched in granulosa and theca cells at diestrus (**B**).

## Potential Limitations

We used online cell type identification databases and prior knowledge to define labels for the clusters in the scRNAseq and snRNAseq datasets. However, it is important to note that the RO clusters have not yet been spatially validated, and thus labels may be inaccurate. In addition, the *Pax8rtTA; TreH2BGFP* is more highly expressed in the cells of the EOR and CR than in the IOR, so it’s likely that we have very little representation of the IOR in our datasets.

## Conclusions and Future directions

The datasets described here provide a wealth of information on the transcriptome of all regions of the rete ovarii and surrounding tissue in fetal and adult mice. This work integrates well with recent efforts to better understand the origin and progenitor function of the intraovarian region of the RO in fetal mice and humans (9,11,12). To the best of our knowledge, these are the first transcriptomic insights into the adult RO and the extraovarian regions of the RO. We are actively working on spatial validation of the data presented here, with the goal of identifying candidate genes that may lead us to functions of the RO. Importantly, we hope these datasets become important resources in the field of ovarian biology that may provide novel candidates for the investigation of idiopathic female subfertility.

## Code Availability

All quality control analyses were performed using FastQC (http://www.bioinformatics.babraham.ac.uk/projects/fastqc/). Bulk RNAseq analysis was performed using Salmon (https://combine-lab.github.io/salmon/) and DESeq2 (https://github.com/thelovelab/DESeq2).

Single-cell and single-nucleus analysis was performed using Cell Ranger (downloaded from 10x genomics) and Seurat (https://satijalab.org/seurat/).

## Acknowledgements

The authors would like to thank Dr. Nicolas Devos from the Duke Center for Genomic and Computational Biology Sequencing and Genomic Technologies core facility for sequencing resources and assistance; members of the Duke Flow Cytometry facility for helpful guidance and FAC-sorting; and the Duke Office of Information Technology and the Duke Compute Cluster and team for the assistance and computational power to perform the initial sequencing analyses reported here. We are grateful to all members of the Capel laboratory, the McKey laboratory and the Roberson laboratory for daily inspiration and helpful discussions and suggestions on the work presented here.

## Author contributions

DNA, YX, DTP and JM carried out the experiments; DNA, RO, and JM carried out the data processing and data analysis, RO formatted and deposited the data in appropriate repositories and designed the shiny app to query the data; DNA, BC and JM designed the experiments; DNA, BC and JM drafted, revised and edited the manuscript; BC and JM acquired funding for the project.

